# Single-cell analysis reveals cellular heterogeneity and molecular determinants of hypothalamic leptin-receptor cells

**DOI:** 10.1101/2020.07.23.217729

**Authors:** N. Kakava-Georgiadou, J.F. Severens, A.M. Jørgensen, I. Stoltenborg, K.M. Garner, M.C.M Luijendijk, V. Drkelic, R. van Dijk, S.L. Dickson, T.H. Pers, O. Basak, R.A.H. Adan

## Abstract

Hypothalamic nuclei which regulate homeostatic functions express leptin receptor (LepR), the primary target of the satiety hormone leptin. Single-cell RNA sequencing (scRNA-seq) has facilitated the discovery of a variety of hypothalamic cell types. However, low abundance of LepR transcripts prevented further characterization of LepR cells. Therefore, we perform scRNA-seq on isolated LepR cells and identify eight neuronal clusters, including three uncharacterized Trh-expressing populations as well as 17 non-neuronal populations including tanycytes, oligodendrocytes and endothelial cells. Food restriction had a major impact on Agrp neurons and changed the expression of obesity-associated genes. Multiple cell clusters were enriched for GWAS signals of obesity. We further explored changes in the gene regulatory landscape of LepR cell types. We thus reveal the molecular signature of distinct populations with diverse neurochemical profiles, which will aid efforts to illuminate the multi-functional nature of leptin’s action in the hypothalamus.

## Introduction

Leptin is a hormone predominantly produced by white adipose tissue and secreted into the blood circulation^1^. Leptin regulates metabolism and appetite by inhibiting food intake, lowering body weight and increasing metabolic rate^2^. Circulating leptin levels are proportional to body fat amount and body mass index (BMI)^3^. Leptin also responds to acute changes in energy balance: fasting decreases and feeding increases leptin levels^4^. Obese people have high circulating leptin levels suggesting decreased sensitivity to leptin^3,4^, which is referred to as leptin resistance.

The hypothalamus plays a major role in the effects of leptin on energy balance, with strong leptin receptor (LepR) expression in the Arcuate nucleus (Arc). The two most well studied LepR neurons in the Arc are the orexigenic AGRP/NPY and the anorexigenic POMC/CART neurons. Leptin inhibits Agrp and stimulates Pomc neurons^5–7^, while Agrp neurons are the most sensitive hypothalamic cell population to energy deficit^8–10^.

LepR is also expressed in less defined neurons in the ventromedial hypothalamus (VMH), dorsomedial hypothalamus (DMH), preoptic area (PO), pre-mammillary nucleus (PMv), lateral hypothalamus (LH) and weakly in the paraventricular nucleus (PVN)^11–13^, that contribute to leptin’s effects on homeostatic functions^14–16^. Besides neuronal cells, LepR expression has also been reported in astrocytes^17,18^, tanycytes^19^, endothelial cells^20^, microglia and macrophages^21^, oligodendrocytes^22^ and vascular leptomeningeal cells (VLMCs)^23^.

Thus, the hypothalamic LepR population of cells is highly diverse, but a full description of the diversity of their transcriptomes remains unresolved. Unravelling the transcriptome of hypothalamic leptin receptor expressing cells helps to identify cell-specific targets to treat obesity. In the mouse hypothalamus, single-cell RNA-seq has provided valuable information about the molecular profile of hypothalamic cell types and new neuronal and non-neuronal subtypes^9,10,24^. However, in hypothalamic scRNAseq datasets it is difficult to identify LepR cells due to their scarcity and the low abundance of LepR transcripts.

To overcome these hurdles, we took a biased approach to identify all LepR-expressing cells in the hypothalamus. We used a LepR-Cre mouse crossed with a mouse expressing the Cre-dependent fluorescent protein tdTomato to specifically isolate hypothalamic LepRcells using FACS and sequence their transcriptome at single-cell resolution. We identify new LepR subtypes, define their molecular signatures, active gene regulatory networks and their enrichment in GWAS Body Mass index signals. Furthermore, we show that fasting induces major transcriptional changes in Agrp LepR neurons. Overall, our dataset provides insight into the molecular makeup of hypothalamic LepR cells and a source for future studies.

## Results

To characterize LepR^+^ hypothalamic cells, we optimized isolation of intact single cells from the hypothalamus of LepR-tdTomato mice (see **Methods**). The tdTomato-expressing cells from fed or 24h-fasted animals were FACS-sorted based on tdTomato fluorescence (**Fig. 1a, S1a**) followed by single-cell RNA sequencing (scRNAseq, see **Methods**).

**Figure 1.**
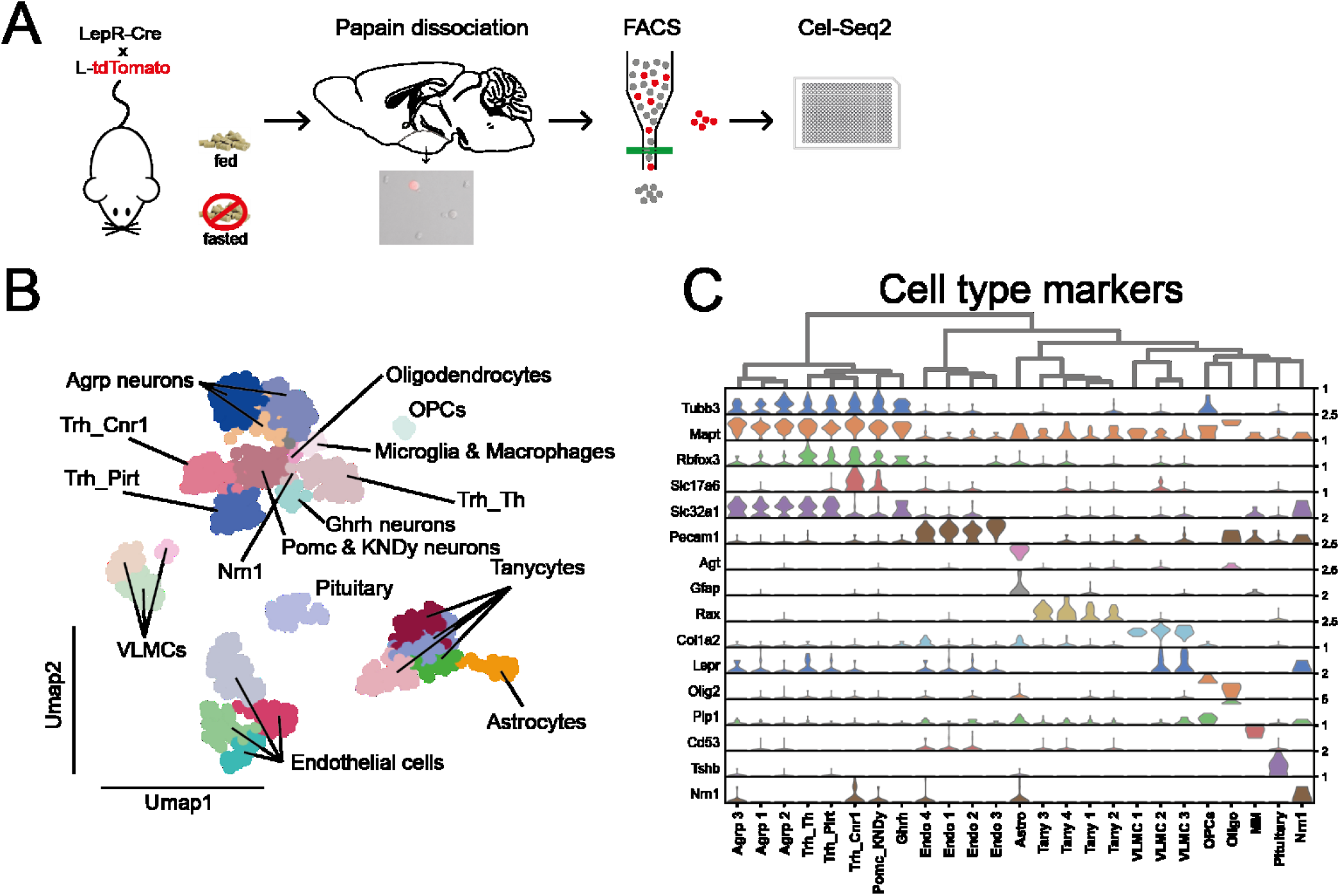
scRNAseq on hypothalamic LepR cells reveals 25 clusters. **A.** Hypothalami of fed and 24-hour fasted LepR-Cre x L-tdTomato mice were dissected, dissociated with papain and FACS-sorted into 384-well plates, where the Cel-Seq2 protocol was performed. **B.** 25 clusters of hypothalamic LepR cells annotated on a UMAP after clustering analysis. **C.** Violin plots of pan marker genes used to define which main cell types the clusters represent. The maximum normalized count of each pan marker gene is presented on the right. Endo: Endothelial, Astro: Astrocytes, Tany: Tanycytes, VLMC: Vascular leptomeningeal cells, MM: Microglia and Macrophages, Oligo: Oligodendrocytes, OPCs: Oligodendrocyte progenitor cells.

A total of 1,288 tdTomato^+^ cells were sorted in four plates/batches (**Fig. S1b**) and were sequenced with Cel-Seq2^25^. Of all tdTomato cells, 1,048 cells were from fed animals (“fed cells”) and 240 cells from fasted animals (“fasted cells”) and were subsequently used for analysis using Scanpy^26^. Cells containing less than 2,000 genes and with more than 50% of the reads belonging to ERCC spike-in RNA (indicative of poor caption of mRNA from cells) or with more than 15% mitochondrial gene expression (indicative of RNA degradation and poor cell viability) were excluded (**Fig.S1c**). These steps left 824 cells for analysis, 692 cells from the fed condition and 132 cells from the fasted condition (**Fig. S1b**), with a high-quality dataset containing a median of 12,082 unique counts and 4,756 genes per cell. To overcome the effect of sequencing depth between neurons and non-neuronal cells in clustering, we used downsampling prior to the analysis (see **Methods** for details).

Unsupervised clustering using the graph—based Leiden algorithm identified a total of 25 clusters (**Fig. 1b**), which were further assigned to cell types based on expression of established cell type markers (**Fig. 1c**). Hierarchical clustering indicated a clear separation of neuronal and non-neuronal clusters (**Fig. S2a**). Eight neuronal clusters with 373 cells were identified based on the expression of both *Tubb3* (class III β-tubulin) and *Mapt*, whereas 451 cells belonged to 17 non-neuronal clusters, as defined based on the absence of both markers (**Fig. 1c**). We also identified a cluster of pituitary cells using *Tshb* as a marker and excluded it from further analysis. The high complexity of our scRNAseq dataset allows us to detect rarely expressed genes in specific cell types, which are notoriously hard to detect with single cell methods. As expected, *Lepr* mRNA was not identified in all cells, due to the low abundance of LepR transcripts, but it was most prominently expressed in Agrp, Trh_Th and Vascular leptomeningeal cells (VLMC) clusters (**Fig. S2b**).

### Neuronal cells

Eight neuronal (*Tubb3*^+^*Mapt*^+^) clusters were identified and assigned to subtypes based on known or novel marker genes (**Fig. 1b, c** and **Fig. 2a, b**). Based on established marker genes, the classification revealed three **Agrp** neuron clusters with high *Agrp* and *Npy* expression, one **Pomc_KNDy** neuron cluster with high expression of *Pomc*, *Cartpt*, *Kiss1*, *Pdyn* and *Tac2* as well as one **Ghrh** neuron cluster with high *Ghrh* and *Gal* expression. The remaining three clusters could not be assigned based on unique genes; therefore, they were designated based on expression of *Trh* as well as genes that strongly marked the cluster: **Trh_Pirt**, **Trh_Cnr1** and **Trh_Th (Fig. 1c, 2a, 2b)**.

**Figure 2.**
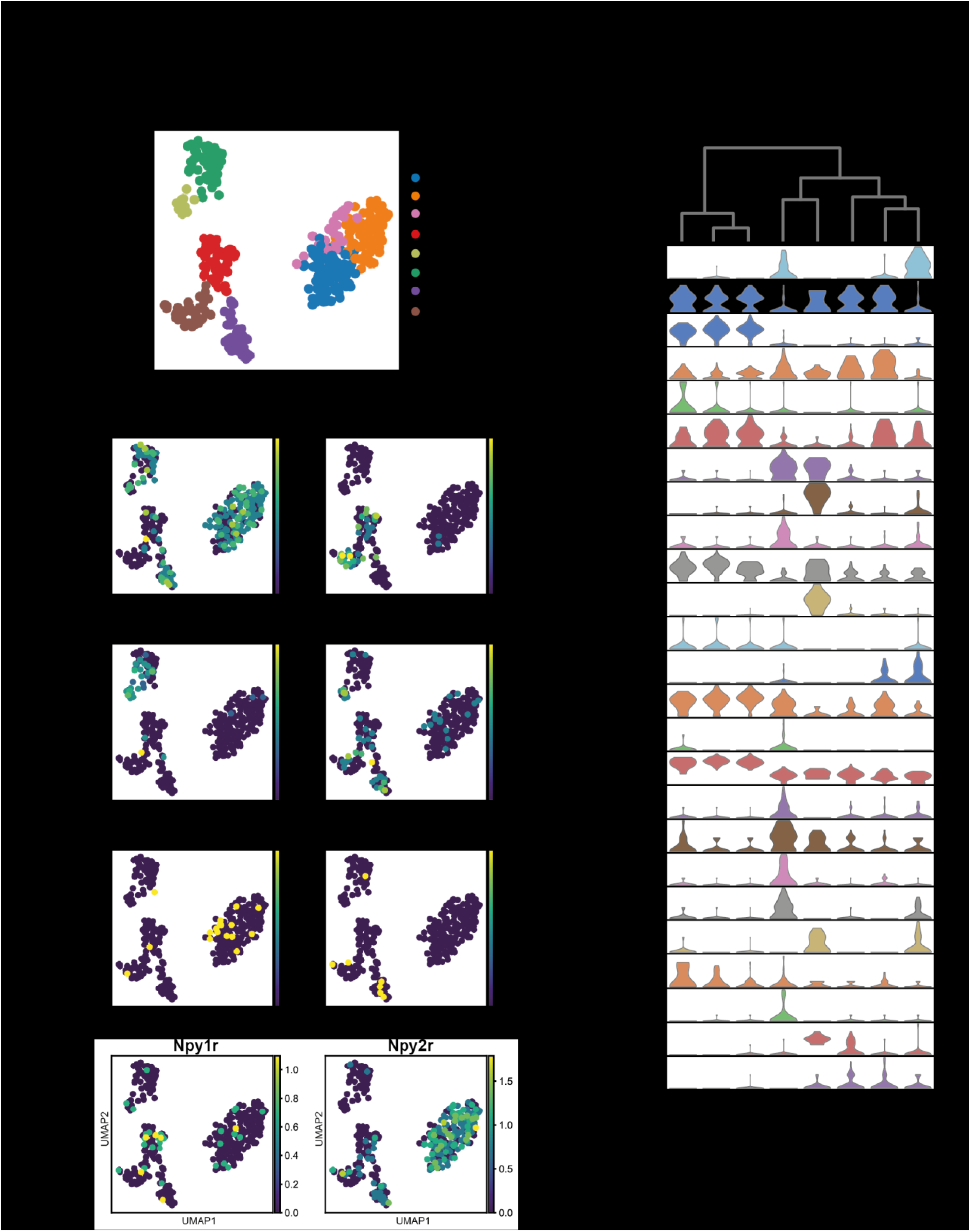
Overview of the eight neuronal LepR clusters in the hypothalamus. **A.** 8 neuronal clusters of LepR cells annotated on a UMAP plot. **B.** Violin plots of expression of genes used to characterize the neuronal clusters, including GABAergic marker *Slc32a1*, glutamatergic marker *Slc17a6*, various neuropeptides and receptors. The maximum normalized count of each gene is presented on the right. **C-F.** UMAP plots showing expression level of genes in neuronal clusters. The expression level is color-coded.

Based on the expression of *Slc32a1* (which codes for VGAT) and *Slc17a6* (which codes for VGLUT2) (**Fig. 2b, 2c**), **Agrp**, **Ghrh** and **Trh_Th** clusters were GABAergic, while the **Pomc_KNDy** neuron cluster contained both GABAergic and glutamatergic cells in line with the literature^27–29^. The **Trh_Pirt** and **Trh_Cnr1** clusters were comprised of mostly GABAergic cells with a few glutamatergic cells.

The dopamine marker *Th* was most prominently expressed in the **Ghrh** and **Trh_Th** clusters (**Fig. 2d – left**). The dopaminergic marker *Ddc* was also expressed in the **Ghrh** cluster but mostly marked the **Trh_Pirt** and **Trh_Cnr1** clusters (**Fig. 2d – right**). Besides the **Pomc_KNDy** cluster, high expression of *Cartpt* was also found in the **Trh_Th** and **Trh_Pirt** clusters (**Fig. 2b**). In line with what is known about LepR neurons, neither *Hcrt* nor *Pmch* were expressed in our dataset (not shown). Few cells expressing *Crhr2* were found in clusters **Trh_Pirt**, **Trh_Cnr1** and **Trh_Th**, while most cells with *Crhr1* expression were found in **Agrp** clusters (**Fig. 2e**).

Moreover, the serotonin receptor gene *Htr2c*, a receptor widely distributed in hypothalamic LepR cells and a drug target of the obesity drug Lorcaserin^30,31^, was significantly enriched in the **Trh_Cnr1** cluster, with high expression in the **Trh_Pirt** cluster, while it was also detected in a limited number of **Pomc_KNDy** cells (**Fig. 2b**), in which serotonin 2c receptors have an established role in energy homeostasis^32,33^. Interestingly, Campbell et al.^10^ observed high expression of *Lepr* in a distinct POMC subset that was negative for *Htr2c*. *Esr1* (Estrogen receptor 1) has an important role in metabolic regulation^34^ and was highly expressed in the POMC_KNDy and Ghrh clusters (**Fig. 2b**). A full list of expression profiles of neuropeptides and their receptors is provided in the supplementary figures (**Fig. S3a, b**).

Next, we looked into the molecular signatures of specific neuronal clusters. **Agrp** neurons divided into 3 clusters, which all showed high expression of *Ghr* (growth hormone receptor), *Crhr1* (corticotropin-releasing factor receptor 1), *Irs4* (Insulin receptor substrate 4), *Ghsr* (ghrelin receptor), *Egr1* (early growth response protein 1) and *Sst* (somatostatin) with higher enrichment in Agrp cluster 3 (**Fig. 2b**). As expected, the *Npy2r* (neuropeptide Y receptor 2), a presynaptic autoreceptor, was expressed in almost all Agrp neurons^35^, along with cells in the **Pomc_KNDy**, **Trh_Pirt** and **Trh_Cnr1** clusters, whereas *Npy1r* (neuropeptide Y receptor 1) was mostly expressed in **Pomc_KNDy** neurons, along with other cell types (**Fig. 2f**).

The **Pomc_KNDy** cluster displayed high expression of *Prlr* (Prolactin receptor), *Pgr* (Progesterone receptor) and *Gfra1* (GDNF family receptor alpha-1) (**Fig. 2b**).The **Ghrh** neuron cluster showed enrichment of *Gal* (Galanin) and *Reln* (Reelin), both of which were also expressed in lower levels in different subsets of cells in cluster **Trh_Cnr1**(**Fig. 2b**). Most of the LepR GHRH neurons are located in the Arc and half of them engage in leptin-mediated pSTAT3^30^. Somatostatin inhibits whereas leptin stimulates pulsatile GHRH-mediated growth hormone release from the pituitary^36–39^. While Arc GHRH neurons express both Somatostatin receptors (*Sstr1* and *Sstr2*)^40^, in our LepR Ghrh cluster we found expression of solely *Sstr2* (**Fig. S3b**). Even though knock-out of LepR in GHRH neurons does not affect body weight^30^, this LepR population poses exceptional interest for further studies.

Three clusters showed expression of *Trh* (**Fig. 3a**). **Trh_Th** neurons were GABAergic neurons that also showed high expression of *Cartpt* (**Fig. 2b, 2d**), while almost half of them expressed *Th* (**Fig. 2d**). In this cluster, there was high expression of transcription factors *Lhx1*, *Lhx1os* (**Fig. 3a**), being co-expressed in 3 cells (**Fig. 3b**), as well as *Kit* (**Fig. 3a**). Receptors *Ptger3* and *Adra1b*, expressed in the preoptic nucleus (PON) and involved in BAT thermogenesis^41–43^ were also highly expressed (**Fig. 3a, 3b**). Many cells in this cluster also expressed *Cxcl12* (**Fig. 3a**), a chemokine mainly expressed in the PVH, LH and lower in Arc, with its mRNA levels increasing upon high-fat diet in these nuclei^44^.

**Figure 3.**
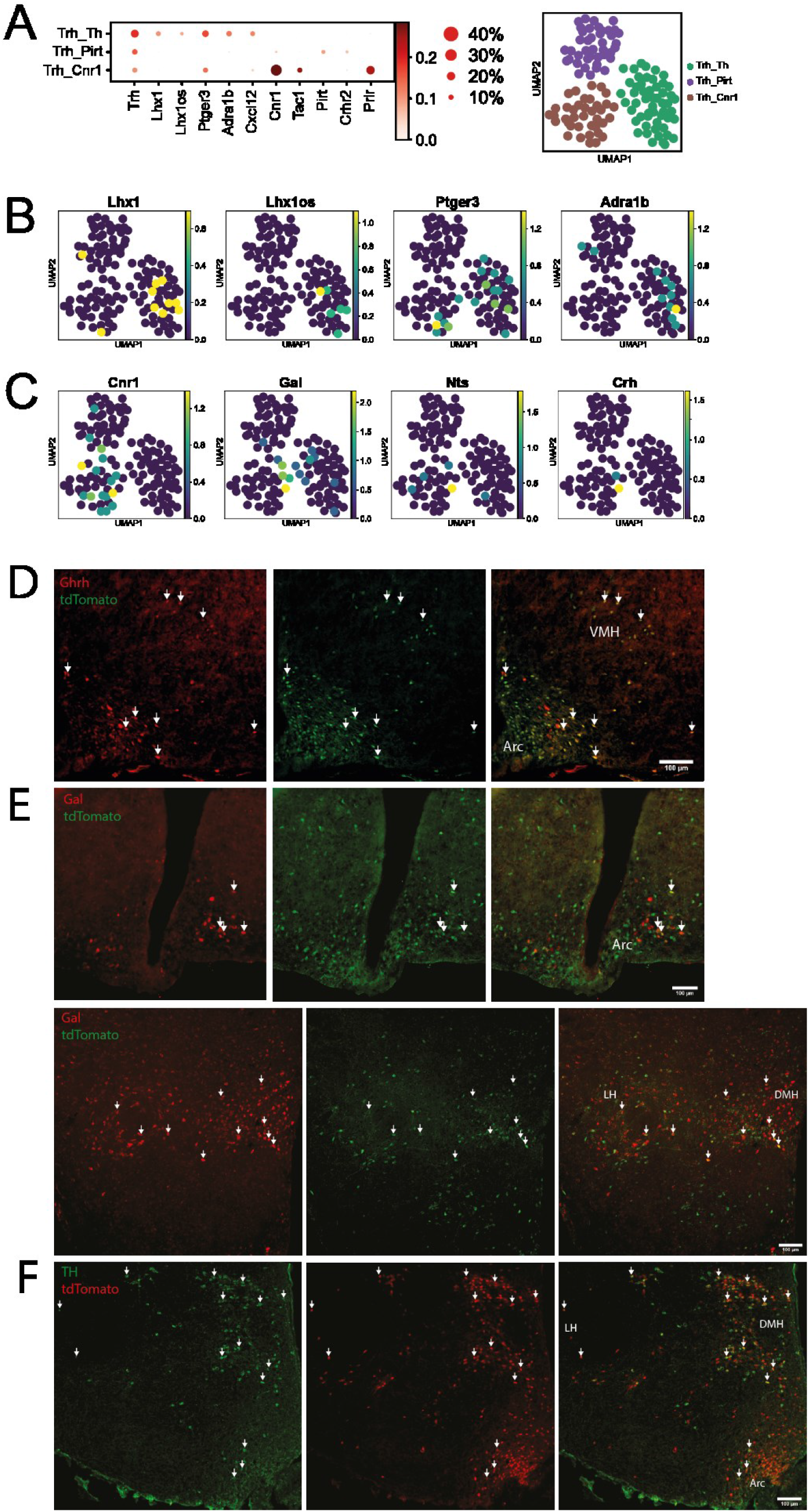
Overview of the 3 novel Trh clusters. **A.** Left: Dot-plot of genes expressed in the three Trh clusters. Color-coding represents expression levels, dot size represents the percentage of cells expressing this gene. Right: Trh neurons were reanalyzed. The three clusters are shown on a UMAP plot. **B-C.** UMAP plots showing expression level of selected genes in Trh clusters. The expression level is color-coded. **D.** Pictures of fixed hypothalamic sections from LepR-Cre x L-tdTomato mice stained with an antibody against tdTomato (green) and fluorescent in situ hybridization (FISH) with anti-sense probe against *Ghrh* (red). **E.** Pictures of fixed hypothalamic sections from LepR-Cre x L-tdTomato mice stained with an antibody against tdTomato (green) and fluorescent in situ hybridization (FISH) with anti-sense probe against *Gal* (red) **F.** Pictures of fixed hypothalamic sections from LepR-Cre x L-tdTomato mice stained with an antibody against TH (green). tdTomato fluorescence appears in red, there was no staining performed against tdTomato. White arrows indicate cells in which co-localization is observed. Scale bars: 100 um. Arc: Arcuate nucleus, VMH: Ventromedial hypothalamus, DMH: Dorsomedial hypothalamus, LH: Lateral hypothalamus

High *Th* expression was observed in both the **Ghrh** and **Trh_Th** clusters, with high expression of *Ghrh* and *Gal* in the former and low or absent expression of these markers in the latter. We therefore investigated the expression patterns of Th, *Ghrh* and *Gal* in LepR neurons in the hypothalamus, by performing IHC and FISH on hypothalamic tissue from LepR-Cre x L-tdTomato mice. *Ghrh* mRNA was found to be co-expressed with tdTomato mostly in the Arc and to a lesser extent in the VMH (**Fig. 3d**), confirming the Arc as a major co-localization site^30^, while *Gal* mRNA co-localized with tdTomato in the Arc (**Fig. 3e - top**), DMH and LH (**Fig. 3e - bottom**). Th signal was found in tdTomato-positive cells in the Arc, DMH and LH (**Fig. 3f**). This supports that Ghrh^+^Gal^+^Th^+^LepR^+^ neurons are located in the Arc, while Trh_Th LepR neurons are a distinct population of neurons potentially located in the Arc, DMH, PVH, PON and LH. Of note, Campbell et al.^10^, 2017 who performed scRNAseq on the Arc and ME, identified a leptin-sensing GABAergic cluster (n11.Trh/Cxcl12) with high expression of *Trh*, *Kit*, *Ptger3* and *Cxcl12*, all genes that were also found in the Trh_Th cluster.

The **Trh_Cnr1** cluster was best defined by high expression of *Cnr1* (cannabinoid receptor 1) (**Fig. 3a, c**), which is strongly linked to feeding behavior^45^, and also displayed high expession of *Prlr* (Prolactin receptor). This cluster also contained small subsets of neurons expressing a diversity of neuropeptides. Few cells in this cluster expressed *Bdnf*, which regulates feeding and energy balance^46–48^. *Cnr1* and *Bdnf* are most strongly expressed in the VMH and weaklier in other hypothalamic nuclei (but not Arc), based on Allen Brain Atlas ISH data and literature^48–50^.

A subset of glutamatergic neurons in the **Trh_Cnr1** cluster showed expression of *Tac1* (Tachykinin 1) (**Fig. 3a**), which has been identified as a LepR neuron subtype marker in the PMv and LH by TRAP-seq and is linked to feeding behaviour and metabolism^51,52^. We also found expression and co-localization between markers *Gal*, *Nts* and *Crh*, established as LepR markers in the LH^51,53^, with one GABAergic cell co-expressing all three (**Fig. 3c**). In our histochemical analysis above, we also find co-expression of *Gal* mRNA with tdTomato in the DMH, besides the LH. This suggests that neurons in this cluster are not centralized in a single hypothalamic nucleus, but are most likely dispersed in the VMH, DMH and LH hypothalamic nuclei.

Neurons in the **Trh_Pirt** cluster were GABAergic and showed relatively high expression of *Cartpt* (**Fig. 2b, d**) in almost all of the cells of the cluster, while few cells express *Crhr2* (**Fig. 2e**). Few cells expressing *Pirt* (Phosphoinositide-interacting protein), with a role in obesity^54^, were exclusively found in this cluster (**Fig.3a**). We also found sparse cells expressing *Prlh* (Prolactin Releasing Hormone), *Pthlh* (parathyroid hormone like hormone), *Retn* (Resistin) and *Nmu* (Neuromedin U) (**not shown**).

### Non-neuronal cells

Seventeen (17) non-neuronal clusters, based on the absence of expression of both *Tubb3* and *Mapt*, were assigned to subtypes based on the expression of established canonical markers. Our dataset is composed of four tanycyte [**Tany 1-4**] (*Rax^+^*), four endothelial [**Endo 1-4**] (*Pecam1^+^*), one astrocyte [**Astro**] (*Agt^+^*), one oligodendrocyte [**Oligo**] (*Olig2^+^,* low *Plp1*), one oligodendrocyte progenitor cell [**OPC**] (*Olig2^+^,* high *Plp1*), one microglia and macrophages [**MM**] (*Cd53^+^*), three vascular leptomeningeal cell [**VLMCs 1-3**] (*Col1a2^+^*) clusters and one unknown cluster strongly marked by *Nrn1* (**Fig. 1b, 1c**).

Tanycyte clusters were almost entirely marked by *Rax*, *Vim, Ppp1r1b*, while almost half of the cells expressed *Nes* and only 1 cell *Gfap*, which is mostly found in α tanycyte subtypes^55,56^. Whereas traditionally tanycytes have been classified in 4 categories (α1, α1,β1 and β2), research on their morphology and molecular signature indicates that tanycytes display great heterogeneity and they exist along a gradient^57^. In line with this, we did not observe a clear-cut expression of subtype-specific markers in each cluster (**Fig. 4a**). **Tany 1 & 2** mostly contained α2 and β1 tanycytes, since they were mostly marked by *Crym* and *Frzb*. **Tany3** was mostly marked by the markers of β1 and β2 tanycytes *Col25a1*, *Adm* and *Cacna2d2*, whereas **Tany4** was marked mostly by α2-marker *Vcan* as well as β2-marker *Cacna2d2. Cd59a* and *Slc17a8*, which mark α1 tanycytes were expressed by very few cells. Moreover, **Tany3** displayed high expression of the markers *Tll1* and *Lrp2,* which codes for Megalin and facilitates leptin transport via the BBB with transcytosis^58,59^ and *Cntfr*, confirming that in the ME there are tanycytes responsive to CTNF^60^.

**Figure 4.**
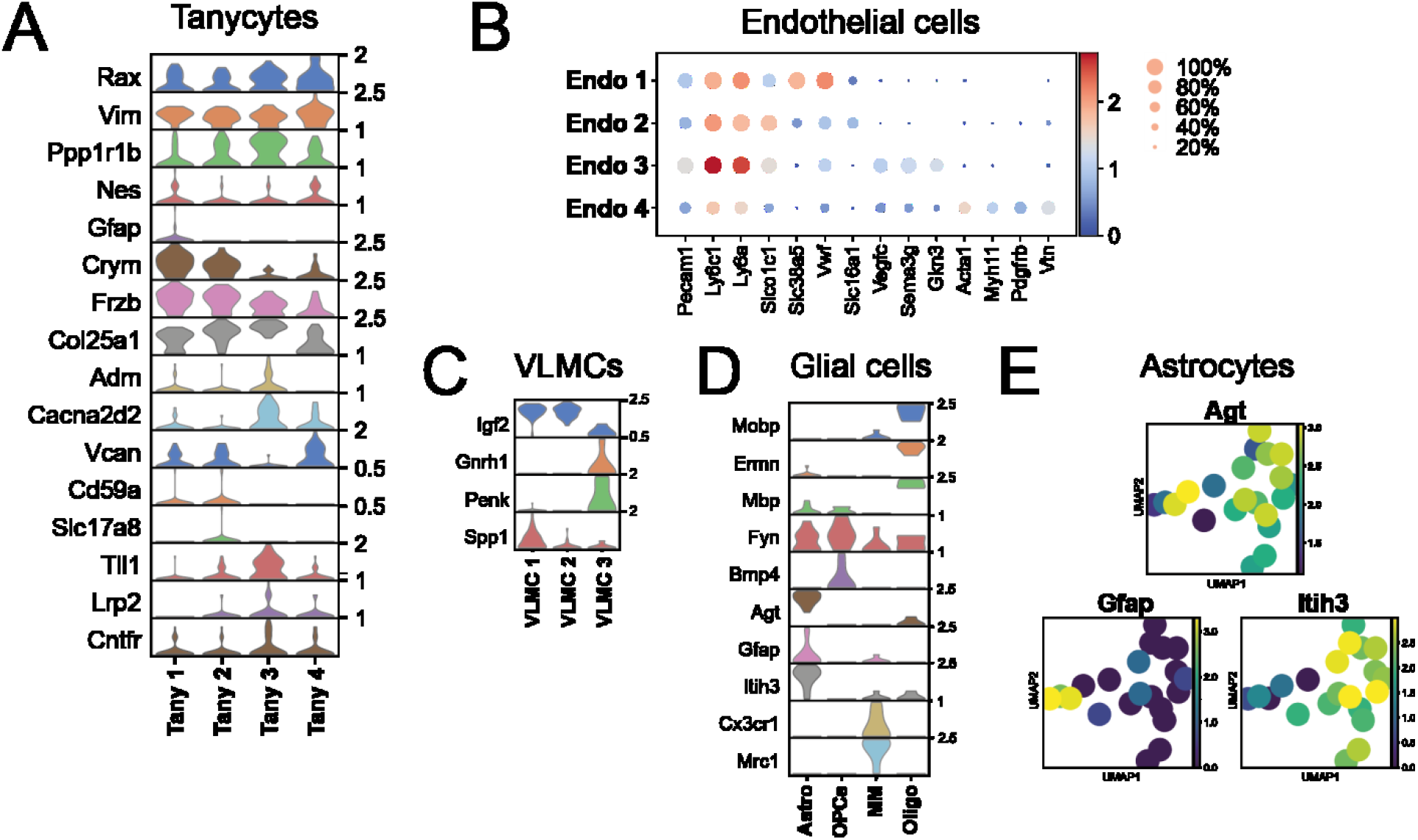
Overview of the non-neuronal clusters of LepR cells in the hypothalamus. **A.** Violin plots showing the expression levels of selected genes in the four tanycyte clusters. **B.** Dot-plot of expression of selected genes in the endothelial clusters. Color-coding represents expression levels and dot size represents the percentage of cells expressing this marker. **C.** Violin plots showing the expression levels of genes in the three VLMC clusters. VLMC: Vascular leptomeningeal cells. **D.** Violin plots showing the expression levels of genes in Glial cell clusters. Astro: Astrocytes, MM: Microglia and Macrophages, Oligo: Oligodendrocytes, OPCs: Oligodendrocyte progenitor cells. **E.** UMAP plots showing expression of astrocytic markers in astrocytes. The expression level is color-coded. In A, C and D, the maximum normalized count of each gene is presented on the right.

We identified 4 clusters of endothelial cells (**Endo 1 - 4**), expressing markers *Pecam1*, *Ly6c1*, *Ly6a* and *Slco1c1*^10,61^, which form a gradient of expression, similarly to tanycytes. **Endo 1** is composed mostly of venous endothelial cells due to the enriched expression of *Slc38a5*and *Vwf*, whereas in **Endo 2***Slc16a1* is highly expressed, which marks capillary cells, and **Endo 3** consists of mostly arterial cells highly enriched in *Vegfc, Sema3g* and*Gkn3*. **Endo 4** seemed to be composed of non-endothelial cells on top of endothelial cells, due to the presence of marker genes of mural cells: *Pdgfrb* and *Vtn* for pericytes and *Acta2, Myh11* for smooth muscle cells^9,10,61^ (**Fig. 4b**).

**VLMC1** shows high expression of *Spp1* (secreted phosphoprotein 1/Osteopontin), which mediates obesity-associated hepatic alterations^62^ and **VLMC1 & 2** show high expression of *Igf2* (Insulin Like Growth Factor 2), whereas **VLMC 3** is marked strongly by *Gnrh1* (Gonadotropin Releasing Hormone 1) and *Penk* (Proenkephalin) (**Fig. 4c**).

Regarding the rest of non-neuronal clusters (**Fig. 4d**), we identified oligodendrocytes, mostly marked by *Mobp* (Myelin Associated Oligodendrocyte Basic Protein), *Ermn* (Ermin) and *Mbp* (Myelin basic protein), while OPCs highly expressed *Fyn* and *Bmp4* (Bone Morphogenetic Protein 4)^9,63^. Astrocytes were strongly marked by *Agt* (Angiotensinogen). *Gfap* was expressed in a small subset of astrocytes, while the rest of the cells in this cluster showed high expression of *Itih3* (**Fig. 4e**), which has 200 fold-higher expression in mouse astrocytes compared to human^64^. In the Microglia and Macrophages cluster we found expression of *Cx3cr1* and *Mrc1*. The expression of LepR in many glial populations pinpoints leptin’s important role in many different functions. Studies have shown leptin’s involvement in cytokine production and activation of microglia and macrophages in the context of neuropathic pain^65,66^, while leptin regulates differentiation and/or myelination of oligodendrocytes^63^.

Several unbiased approaches profiled hypothalamus and detected a varying number of cell types. We questioned whether we could assign LepR cells identified in our study to published datasets (**Fig. S4**). For this, we employed Scibet which uses the E-test to predict similar cell types (see Methods, ^67^). Chen et al profiled the entire hypothalamus^9^. LepR non-neuronal cells were easily mapped to specific non-neuronal clusters from Chen et al. (**Fig. S4a**). For instance, **Tany 1-4** were all identified as “Tanycytes”, while **Endo 1-4** and **VLMC 1-3** were assigned to “Epithelial 1-2”. **Agrp neurons 1-3** were identified as “GABA15”, while **Ghrh** and **Pomc** neurons best matched GABA11 and Glu11, respectively. On the other hand, Trh neurons could not be assigned to a specific cluster, suggesting that they are either rare, share similarities with multiple clusters or could only be clustered upon sequencing at higher depth and cell numbers. Direct comparison of Lepr neurons to hypothalamic cell populations may help us determine which option holds true. Overall, we identified several subgroups of glial and neuronal cells that were originally assigned as a single cell type in other studies as well as three Trh+ cell clusters that were not reported.

### Gene regulatory network of LepR cells

The scRNAseq atlas of LepR cells in the hypothalamus allows us to investigate the molecular mechanisms of their action. Gene regulatory networks (GRNs) are driven by transcription factors that regulate the expression of sets of target genes (gene modules) in a context-dependent manner. We used the pySCENIC algorithm to determine activity of ‘regulons’, groups of genes that are co-expressed with a transcription factor that drives their expression, in single cells^68,69^. We detected 38 regulons that display a spectrum of activity over LepR cells, which we visualized using t-SNE (**Fig. S5a**). Regulon activity recapitulated the distribution of cells types, suggesting that the major determinants of cell types are captured by the GRNs (**Fig. 5a**).

**Figure 5.**
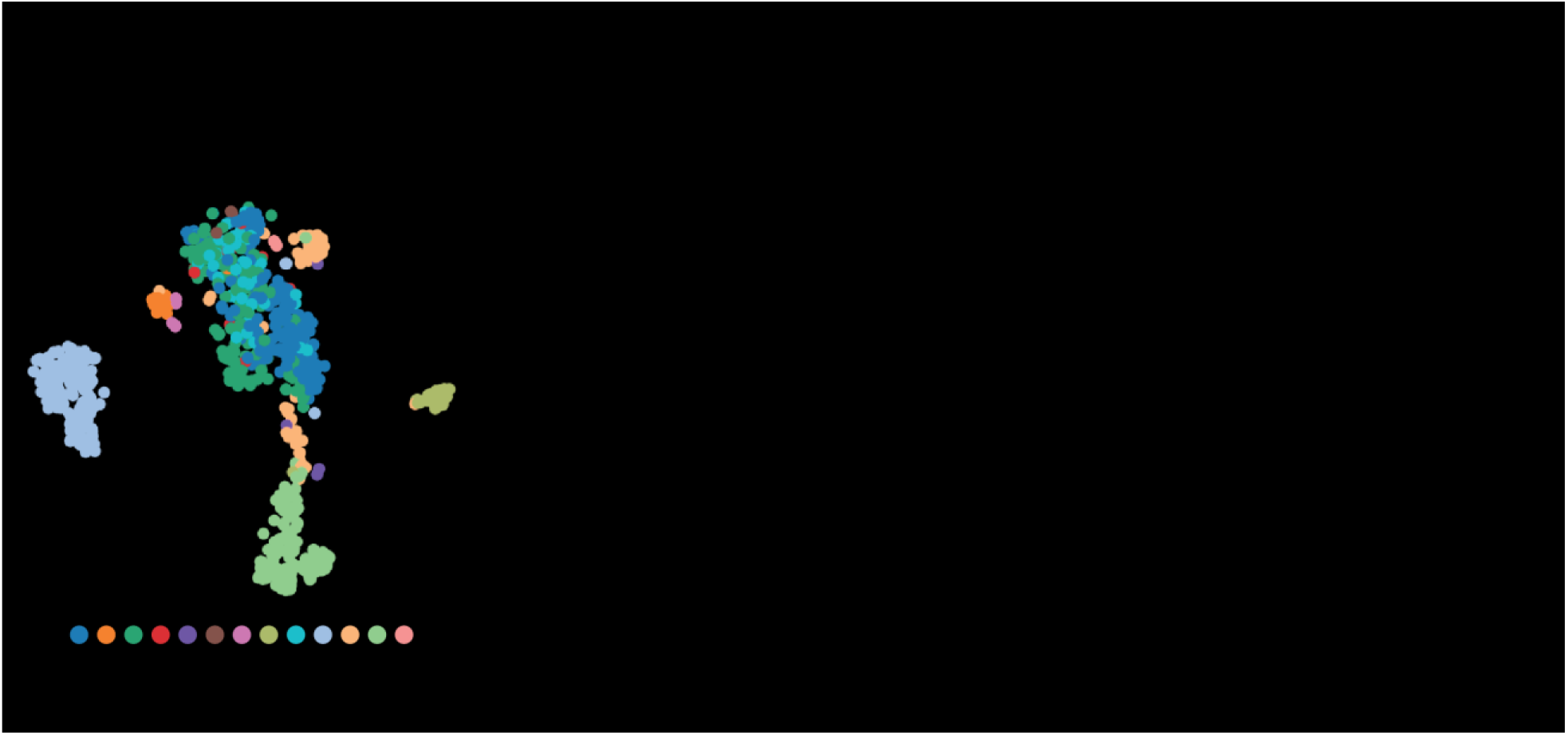
Gene regulatory network (GRN) analysis of LepR cells in the hypothalamus by pySCENIC. **A.** tSNE plot showing cell categories following clustering based on GRN analysis. **B.** Dendrogram representing hierarchical clustering based on regulons identified as active by pySCENIC analysis. The main regulons that are driving discrimination of branches (cell types) are annotated. Endo: Endothelial, Astro: Astrocytes, Tany: Tanycytes, VLMC: Vascular leptomeningeal cells, MM: Microglia and Macrophages, Oligo: Oligodendrocytes, OPCs: Oligodendrocyte progenitor cells.

Next, we performed hierarchical clustering to group cell types using their regulon activity (**Fig. 5b**). Neurons, astroglia and non-neural cells formed clearly distinct branches in the hierarchy, and subgroups of cell types clustered together (**Fig. 5b**). Next, we interrogated the regulation networks that were differentially active between different branches of the LepR cell hierarchy using the Mann-U-Whitney test (**Fig. 5b**, see **Methods**). Npdc1 and Smarcc2 regulons discriminated the neuronal cells from astroglia, which was enriched in Ets1 and Erg regulons. Among the neuronal cell types, higher activity of Fli1 and Bclaf1 further defined Agrp neurons. These analyses provide a rich source of regulon activity (**Fig. S5b**).

### Regulation of LepR-Agrp neurons by fasting

Next, we investigated the transcriptional response of LepR/tdTomato hypothalamic cells to energy deficit (24-hour fasting). Among the clusters, we saw that Agrp neurons responded the most to fasting, with the highest number of genes being differentially expressed (not shown). We cumulated the Agrp clusters to delineate the effect of fasting on LepR cells. Within the Agrp neuron population fasting induced upregulation of *Lepr*, *Agrp* and *Npy* (**Fig. 6a**). Using Wilcoxon rank sum test, we identified 93 genes upregulated and 13 genes downregulated by fasting with log2FC>1.5 and <-1.5 respectively (**sup. Table 2**). Confirming previously identified genes in Agrp neurons^8^, fasting also induced upregulation of *Ghsr*, *Mt1*, *Acvr1c* and *Vgf* (**Fig. 6a**), all of which are involved in obesity and energy balance^70–73^ and downregulation of circadian clock genes *Nr1d2*, *Per3* and *Bhlhe40* (**Fig. 6b**).

**Figure 6.**
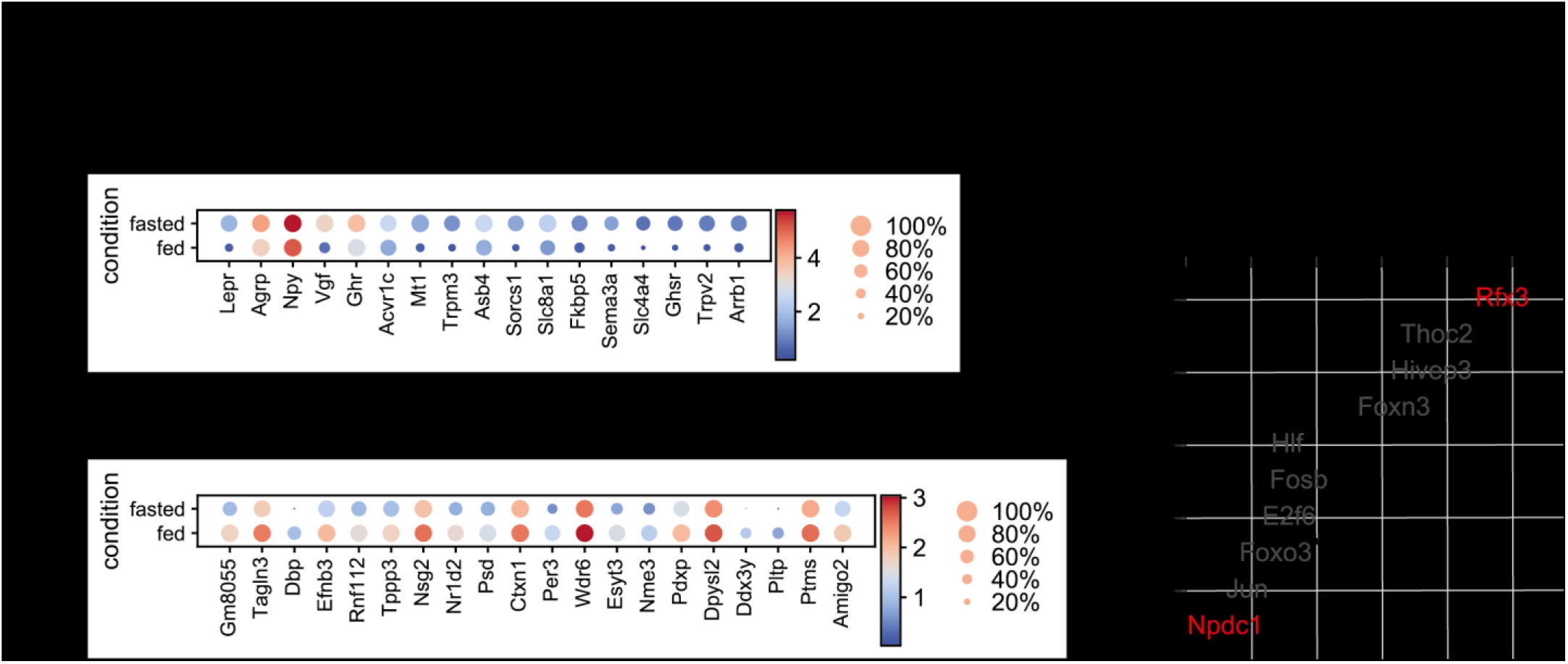
Effect of 24-hour food deprivation on LepR Agrp neurons. **A-B.** Dot-plots of a selection of genes identified to be upregulated and downregulated by fasting in LepR Agrp neurons. Color-coding represents expression levels, dot size represents the percentage of cells expressing this marker. In A, selected genes are shown. Besides *Lepr*, *Agrp* and *Npy* displayed on the left, the rest of the selected genes are ordered from left to right by lowest to highest p value. In B, the top 20 downregulated genes are ordered from left to right by lowest to highest p value. **C** Scatter plot showing how regulon activity is affected by fasting in LepR Agrp neurons, ranked by regulon score. p-adjusted: **<0.001, *<0.05, following Mann-Whitney-Wilcoxon test.

Among the upregulated genes (**Fig. 6a**) we find the transporters *Slc4a4* and *Slc8a1*, the receptors *Ghr* with a role in hepatic glucose production^74^, *Sorcs1* and *Trpm3* with a role in diabetes, energy balance and temperature sensing^75–77^ as well as the cation channel *Trpv2* which regulates BAT thermogenesis^78^. Other upregulated genes that have a relation to obesity and energy balance are *Sema3a*, *Arrb1*, *Asb4* and *Fkbp5*^79–83^.

Next, we asked whether we can identify the GRNs involved in response to fasting in Agrp neurons. We reanalyzed the cells from Agrp clusters, which had a median of ~32000 reads and ~9000 genes per cell, and identified 10 regulons using pySCENIC (**Fig. S6a**, see **Methods**). Comparison of fed and fasted Agrp neurons revealed that Rfx3 (p=0.000012) and Npdc1 (p=0.00016) were the most significantly up- and down-regulated regulons upon fasting, respectively (**Fig. 6c**). While Rfx3 (p=0.0038) mRNA expression was higher in fasted cells, Npdc1 (p=0.756) mRNA expression was not significantly different between groups, suggesting that the activity of the latter is controlled post-transcriptionally (**Fig. S6b**).

Regarding the target genes of Rfx3 and Npdc1 (**sup. Table 2**), we explored their association with energy balance. A target of both Rfx3 and Npdc1, *Mapk10* (also known as JNK3), with a role in in energy balance and insulin resistance^84–87^, protects against excessive adipocity^88^. Its selective deficiency from LepR Agrp neurons causes hyperhagia^89^. A downstream target of Rfx3 is the BDNF receptor *Ntrk2* (also known as TrkB) which maintains negative energy balance mainly by suppressing feeding^90–94^. Significant hits also were the Npdc1 targets *Clstn3* and *Faim2*. *Clstn3* is a key gene that mediates the neuro-adipose junction formation or remodeling in white adipose tissue browning and the BAT activation process^95–97^, while *Faim2* is a GWAS gene associated with obesity and its expression is increased in the Arc upon high-fat diet^98–100^. These results further demonstrate the use of our dataset for delineation of the molecular mechanisms governing LepR neuronal activity.

### LepR cell enrichment for BMI GWAS genes

We next asked whether genes specifically expressed within the LepR cell types enriched for genes implicated by genome-wide association study data for obesity (GWAS). Towards that end we first used CELLEX to identify specifically expressed genes for each cell type and then applied CELLECT^101^ to compute the enrichment of GWAS signal for each set of cell type-specific genes. As genetic association data for obesity we relied on UK Biobank-based GWAS summary statistics for body-mass index (BMI), a commonly used proxy phenotype for obesity^102^. CELLECT identified three significantly enriched cell types, namely Trh_Cnr1, Trh_Pirt and Pomc_KNDy neurons (Bonferroni P-value threshold, P<0.05/25; **Fig. 7**). Together, these analyses confirm the importance of POMC neurons in obesity and reveal the novel LepR clusters Trh_Cnr1 and Trh_Pirt as potential key important players.

**Figure 7.**
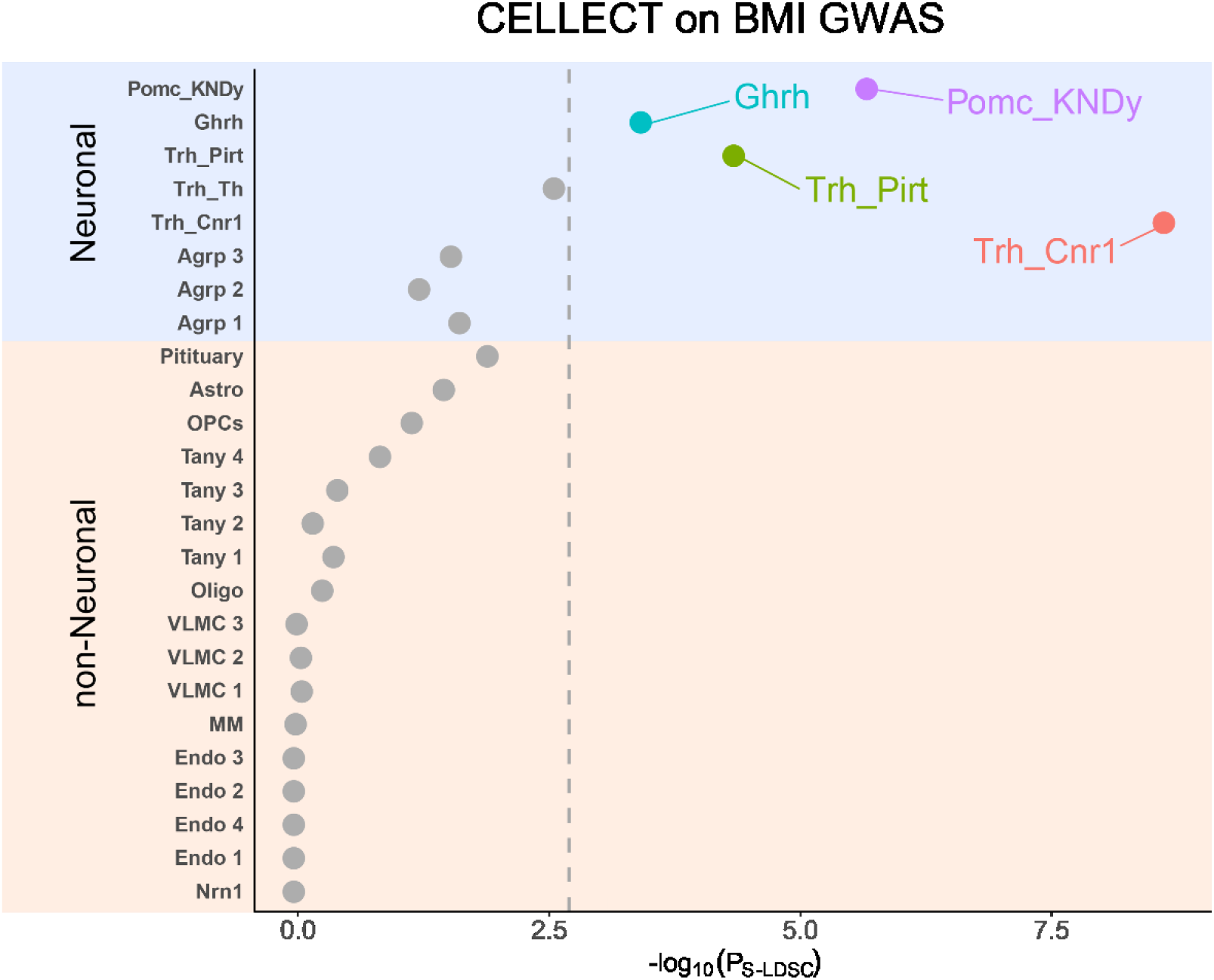
Scatterplot of the CELLECT results. Dashed vertical line marks the Bonferroni threshold for significant *p*-values. The y-axis shows both clustering names and overall cell-type, and the x-axis shows the p-values. Only cell-types that are significant are shown in color Bonferroni P-value threshold (P<0.05/25). S-LDSC, stratified-linkage disequilibrium score regression. Endo: Endothelial, Astro: Astrocytes, Tany: Tanycytes, VLMC: Vascular leptomeningeal cells, MM: Microglia and Macrophages, Oligo: Oligodendrocytes, OPCs: Oligodendrocyte progenitor cells.

## Discussion

We here identified the transcriptome of hypothalamic LepR cells using scRNAseq. Most available scRNAseq datasets focus on one of the hypothalamic subregions and undertake an unbiased approach to identify all cell populations. Instead, we dissociated the whole hypothalamus of LepR-Cre x L-tdTomato mice and focused solely on the leptin receptor-expressing cells. High sequencing depth combined with FACS sorting of tdTomato-positive cells facilitated the identification of twenty-five distinct cell clusters, including eight neuronal and seventeen non-neuronal.

Overall, our dataset highlights the importance of the arcuate nucleus as the primary action site of leptin, with most of the neuronal LepR-expressing cells in this dataset belonging to at least six distinct neuronal populations localized in the Arc as well as tanycyte subtypes which are in close contact to the Arc and ME. Out of the eight neuronal clusters identified, five (Agrp 1-3, Ghrh and Pomc/KNDy) have already been identified as LepR subpopulations, while three (Trh_Th, Trh_Pirt and Trh_Cnr1) are novel. Even though LepR has been identified in Ghrh neurons before^30^, here we show that all neurons in this cluster also express *Th* and *Gal*, revealing a distinct LepR-Arc population with a unique neurochemical profile.

The three newly-identified neuronal clusters displayed *Trh* expression (Trh_Th, Trh_Cnr1, Trh_Pirt) and they appeared to be quite heterogeneous. The enriched genes in these clusters localize in many hypothalamic nuclei based on Allen Brain Atlas ISH data and literature. The **Trh_Th** cluster, with high expression of *Trh*, *Th* and *Cartpt* is highly similar to the Arc leptin-sensing GABAergic Trh/Cxcl12 cluster identified by Campbell et al. Therefore, Trh_Th represents another distinct LepR population in the Arc, even though we cannot exclude the fact that Trh_Th neurons could also be localized in other hypothalamic regions (DMH, PVH, PON, LH), due the expression of receptors *Ptger3*, *Adra1b* and the chemokine *Cxcl12*.

In the **Trh_Cnr1** we identified cells expressing Bdnf, as well as LepR markers of the LH *Nts*, *Gal*, *Tac1* and *Crh*. Remarkably, BDNF neurons in the DMH and VMH do not engage into pSTAT3 signaling upon leptin administration, even though *Bdnf* mRNA is increased^103,104^. While it is known that *Gal* and *Nts* are co-expressed in LepR neurons in the LH^53^, we also found evidence that *Crh* also co-localizes with this neuronal cell type. The expression profiles of this cluster’s enriched genes support that the majority of the neurons are located in the VMH, DMH and LH, skipping the Arc.

The **Trh_Pirt** cluster displayed high expression of *Cartpt*, *Crhr2* and *Pirt*, with the two latter genes being expressed mostly in Arc and VMH. CRFR2 in the VMH has been shown to be critical in feeding and regulation of lipid metabolism in mice^105^. Pirt-KO female mice display an increased susceptibility to develop obesity and glucose intolerance^54^. While on average the cells we analyzed were deeply sequenced, it is likely that sequencing of a higher number of cells may lead to identification of sub-clusters of the cell types we have identified.

Regarding non-neuronal clusters, we identified LepR populations that are important for leptin transport across the BBB (tanycytes, endothelial cells & VLMCs) as well as other glial populations. Tanycyte clusters contained mostly α2- and β1-and β2-tanycytes. The presence of leptin receptor in other non-neuronal cell types, highlights that LepR is not only crucial for leptin transport, but also exerts other functions. Leptin signaling in hypothalamic astrocytes is important for synaptic plasticity and neuroendocrine control of feeding by leptin^18^. Moreover, leptin receptor expression on other glial cells (microglia, macrophages, oligodendrocytes and OPCs) suggests that leptin has a neuroprotective role and it is involved in remyelination, as suggested in literature^21^.

With SCENIC analysis we detected 38 active gene regulatory networks which recapitulated the distribution of cell types among clusters and identified regulons that discriminated cell types from each other, such as Fli1 and Bclaf1 which determined Agrp neurons. Agrp neurons was the population that responded the most to fasting, by increasing the expression hormone receptors involved in energy balance and temperature sensing (*Ghr*, *Ghsr, Sorcs1, Trpm3*) and decreasing the expression of circadian clock genes (*Nr1d2*, *Per3*, *Bhlhe40*), suggesting that fasting makes these neurons more sensitive to changes in feeding status. SCENIC analysis identified Rfx3 and Npdc1 as the regulons that were mostly up- and downregulated by fasting respectively. Their downstream targets *Mapk10*, *Ntrk2*, *Clstn3* and *Faim2* have high association with energy balance and metabolic diseases. Finally, our BMI GWAS enrichment analysis results study confirm the importance of POMC neurons in regulation of energy balance and furthermore they suggest a role for the novel clusters Trh_Cnr1 and Trh_Pirt in regulating body weight.

Thus, the scRNAseq data we provide reveal the heterogeneity of LepR cells in the hypothalamus, including multiple known as well as novel neuronal, astroglial and non-neural cell types. We show that among these LepR cell types, Agrp neurons are the main responders to fasting. We further demonstrate that our dataset can be used to identify the major gene regulatory networks in homeostasis as well as in response to fasting. Overall, we provide a rich source for future studies that dive into the specific roles LepR cell types as well as the molecular circuitry controlling their function.

## Acknowledgements

Part of this work was funded by an NWO Gravitation grant: BRAINSCAPES: A Roadmap from Neurogenetics to Neurobiology: 024.004.012.

## Methods

### Animals

Adult 2-3 month-old ObRb-IRES-Cre mice (B6.129(Cg)-Leprtm2(cre)Rck/J) reported as LepR-Crecrossed with Rosa-CAG-LSL-tdTomato-WPRE::DNeo (008320 007914, Jackson laboratories, Bar Harbor, ME, US) reported as L-tdTomatoon a C57Bl/6J background were used. Mice were housed socially and kept under a 12:12 hr light-dark cycle with lights off at 19:00. Mice were kept at room temperature (21 ± 2°C) and 40-60% of humidity conditions. They were fed with standard chow (Special Diet Service, Essex, UK) and tap water ad libitum. For the 24 hours of fasting protocol, chow was completely removed 24 hours before being killed. Ad libitum mice were male and female and the mice from the fasting protocol were all female. All experiments were approved by the Ani-mal Ethics Committee of Utrecht University and conducted in agreement with Dutch laws (Wet op de Dierproeven, 1996; revised 2014) and European regulations (Guideline 86/609/EEC; Di-rective 2010/63/EU).

### Hypothalamus dissection and single-cell dissociation

In pilot experiments, we optimized dissociation efficiency to obtain viable cells using various conditions including papain, trypsin and accutase A enzymes as well as the use of trehalose, artificial cerebrospinal fluid and several commercially available media. We found that papain was the most optimal enzyme to dissociate hypothalamic tissue with viable tdTomato^+^ cells, and generated an optimized protocol. Mice were killed between 09:00 and 12:00 by manual decapitation after isoflurane anesthesia. Brains were rapidly removed and saved in cold HABG (50 mL Hibernate A without calcium (Brainbits, Cologne, Germany) supplemented with 1 mL B27 and 125 μL L-Glutamine). Hypothalami were microdissected and cut into 1-2 mm thick pieces. Tissue from 2-3 hypothalamiwas pooled into 1-1.5 mL of HABG containing papain (20 u/mL, Worthington Biochemical, Lakewood, NJ, US). Tissue was broken into smaller pieces by pipetting 2-3 times with a P1000 pipet followed by a fire-polished glass pipette with a wide opening. Tubes were incubated for 15’ at 37 degrees, whilst being agitated at 160 rpm. DNAse I (1,000 Kunitz units/mL Worthington Biochemical, Lakewood, NJ, US) was added and tubes were incubated for 15’ at 37 degrees, whilst being agitated at 160 rpm. The samples were chilled on ice the following were added: DNAse I (2,000 Kunitz units/mL), BSA (1.25 μg/mL), FBS (5%) and EDTA (0.5 mM). Next, samples were triturated for 2-3 times with a fire-polished glass pipette with medium opening followed by 2-3 times with a fire-polished glass pipette with a small opening. The triturated sample was passed through a 100um cell strainer. The strained suspension was washed once with 10 mL aCSF(92 mMNaCl, 2.5 mMKCl, 1.2 mM NaH2 PO4, 30 mM NaHCO3, 20 mM HEPES, 25 mM Glucose, 3 mM Sodium Ascorbate, 2 mMThiourea, 3 mM Sodium Pyrurate) supplemented with 10% FBS. Cell suspension was centrifuged at 100 g for 10 minutes at 4 °C. Cell pellet was resuspended in 200-250 μL of collection solution (aCSF containing 150 u/mL DNAse I, 10 μM Rock inhibitor and 10% FBS) and filtered through a 70 μm strainer before sorting.

### Cell sorting

Cells were sorted into 384 well-plates using FACS (BD Biosciences Aria II), after gating for forward and side scatter, selecting singlets and finally the tdTomato+ cells with high fluorescence. Approximately 0.8-0.9% of the total single cell population were defined as tdTomato+ cells (Fig. S1). DAPI was not used for the selection of live (DAPI^−^) cells since pilot experiments showed that tdTomato^+^ cells were almost exclusively live, which was confirmed after RNA-seq analysis by assessing the number of reads and other quality measures (see below). When the plates were not completely filled with tdTomato+ cells, cells from mouse hypothalamic of wild-type (L-tdTomato or C57Bl/6J) mice were used to fill the plates in order to increase the quantity of captured mRNA which helped with production of high-quality libraries. Plates were span at 2000 rpm for 2 min and frozen at −80. In total we processed 7 plates, 4 of which yielded high quality datasets which were used for downstream analysis.

### Library generation, sequencing and alignment

Single-cell RNA sequencing was performed with Cel-seq2 at the Single Cell Discoveries (Utrecht, NL), as described by Hashimshony et al., 2016 and Muraro et al., 2016^25,106^. Plates contained preloaded barcoded poly-T primers that specifically amplify the mRNA and introduce a 6bp unique molecular identifier (UMI), an 8bp cell barcode and a T7 RNA polymerase binding site. In addition, plates were pre-loaded with ERCC spike-ins that allowed quality control of reactions in individual wells. Cells were lysed by heating at 65C for 5 min followed by rapid chilling on ice for 2 min (twice). mRNA was reversed transcribed using Superscript II (ThermoFisher Scientific) and converted into second stranded DNA by DNA polymerase I (ThermoFisher Scientific). Material from each plate was then pooled, cleaned up using Ampure beads (Beckman Coulter, #A63881) and amplified by in vitro transcription overnight using the MEGAscript T7 transcription kit (ThermoFisher Scientific, #AMB1334). Following a second round of clean up, amplified RNA was measured using the Agilent RNA 6000 pico chips (Agilent # 5067-1513), reverse-transcribed using random hexamer primers that introduce Truseq Small RNA kit RP1 primer binding sites (Illumina) and finally converted into DNA libraries using custom rpi primers (RNA PCR Primer Index) adapted from the Truseq Small RNA kit (Illumina)^106^. Following two rounds of Ampure bead clean up and quality control using the Agilent High Sensitivity DNA Kit (5067-4626), libraries were sent for sequencing at the Utrecht Sequencing Facility (USEQ, Life Sciences faculty, Utrecht University) for paired end sequencing (26bp for read 1 and 50bp for read 2) by Nextseq500. 5 plates were combined for a full sequencing run, which yielded ~300M assigned to the libraries.

Alignment of the Celseq2 reads was performed using the mapandgo. Following merging of the reads generated in multiple lanes and removal of the reads that lack Celseq2 barcodes, the data was trimmed using Trimgalore (version 0.4.3^107^) that employs cutadapt^108^ to remove the Illumina library adapters and fastQC^109^ reads with low quality base-calls at the end of the reads. Alignment of the reads to the mm10 reference genome was done STAR (version 2.5.3a^110^), and single-cell libraries were de-multiplexed using the barcoded present in preloaded primer sequences. Following UMI correction, reads were adjusted for the possibility of false assignment of reads from different mRNAs as duplicates due to the low chance of having the same UMI on different mRNA molecules. This was the final count file used for the analysis.

### Computational analysis with Scanpy

We used the Scanpy^26^ pipeline to perform the analysis of our single cell data. We removed cells 2000 genes and with more than 50% of the reads belonging to ERCC spike-in RNA (indicative of poor caption of mRNA from cells) or with more than 15% mitochondrial gene expression (indicative of RNA degradation and poor cell viability) which left 824 cells (out of 1288) for analysis, 692 (out of 1048) cells from the fed condition and 132 cells (out of 240) from the fasted condition. Genes expressed in only one single cell, the *Malat1* gene (due to mapping issues), the ERCC spike-ins and the mitochondrial RNAs were removed from the dataset. This resulted in a high-quality dataset containing a median of 12082 unique counts and 4756 genes per cell.

#### Selection of normalization method

Following log(1+p) transformation and log normalization of the data, we observed that a fourth separate Agrp^+^ cluster was composed exclusively of cells from the fasted condition with higher levels of total read counts. The fraction of ERCC spike-in reads were similar (~9%) between clusters, suggesting that the difference between fed and fasted conditions is not due to a higher amount of total RNA in fasted cells, but due to deeper sequencing (see respective Github page). To test this, we downsampled the reads from each cell to 3000 reads prior to the analysis, which removed the technical differences due to sequencing depth and resulted in fed and fasted cells clustering consistently among the three Agrp clusters. From here on, we used the downsampled dataset and considered the differences as biological. For the analysis of differences between fed and fasted condition in Agrp neurons, we reanalyzed the raw data by downsampling to the median (~39,000 reads per cell). We regressed out the effect of number of genes and the mitochondrial gene expression from both datasets. The data was log-transformed and scaled prior to PCA analysis, which was used for dimensionality reduction using t-distributed stochastic neighbor embedding (t-SNE). Visualization of cell clusters and gene expression was performed using UMAP dimensionality reduction plots, the coordinates of which was used to calculate cell clusters using the Leiden algorithm^111^. Differential gene expression between clusters was performed using the integrated rank_gene_groups() function and the Wilcoxon rank-sum test. Using these marker genes, we annotated cell types. To compare the annotated cell types to the literature, we used the online platform Scibet^67^. The details of clustering, calculation of differential gene expression as well as the genes used for the figures in this paper can be found on https://github.com/neuronur/scLepr.

### Marker gene identification

Spatial expression of the identified genes was investigated from publicly available ISH from the Allen Brain Atlas. Furthermore, for the identified genes PubMed search was applied for literature related to food intake, energy, leptin (receptors) and the hypothalamus. Lowly expressed genes are notoriously hard to detect in scRNAseq, which result in dropouts (zero values in many cells). When expression of rare genes within a cluster is normally distributed, we can consider that the gene is expressed by all cells within the cluster, which our methods stochastically (or randomly) detect.

### Cell type annotation using SciBet

We used the online classification tool (http://scibet.cancer-pku.cn/download_references.html) of SciBet, which employs the E-test to predict similar cell types between the query and a reference. For this, we only included the highly variable genes from the Lepr scRNAseq dataset and used manually annotated cell type names for each cluster determined by the Leiden algorithm. For comparison to Chen et al., the reference dataset provided by the tool was used. For comparison to Campbell et al., we reanalyzed the dataset and exported a data frame of 2000 top highly variable genes and 2000 randomly selected cells. Cell type annotation was provided with the dataset. Details of the preprocessing of the data frames for the analysis can be found at https://github.com/neuronur/scLepr.

### Identification of gene regulatory networks

We used the pySCENIC^68,69^ pipeline to identify gene regulatory networks. In brief, the raw data from all tdTomato^+^ cells or Agrp neurons were used to calculate regulons that contain transcription factors and their targets that are expressed in the same cell using the arboreto package^112^. These regulons were then pruned for targets that lack a binding motif for the respective transcription factor within the 5kb of the transcription start site. The activity of these regulons for each cell were integrated in an Anndata object containing the single cell expression data generated by the scanpy pipeline, as described above. When clustered based on regulon activity, cell types clustered similarly to the analysis based on whole transcriptome. To find out which regulons were driving the branching of cell types upon hierarchical clustering, we used the Wilcoxon rank-sum test. The differences between fed and fasted conditions in Agrp neurons were calculated similarly. A detailed description of the analysis and the code used is available on https://github.com/neuronur/scLepr.

### CELLECT analysis

We used CELLECT v.1.1 ^113^ to identify likely etiologic cell-types underlying obesity. CELLECT takes as input GWAS data and cell-type expression specificity (ES) estimates. The output is a list of prioritized etiologic cell-types for a given complex trait. To generate the ES estimates we used CELLEX v.1.1 ^114^. CELLEX computes robust estimates of ES relying on multiple expression specificity measures (for details see^101^). CELLEX was run using the raw gene expression matrix and the metadata containing clustering information. The resulting cell-type specificity matrix along with the UKBB BMI GWAS ^102^ was used as input for CELLECT which was run with default parameters. For the associations of our dataset to additional GWAS studies, the following resources were used:^115–123^ & Alkes group at the Broad institute and https://www.med.unc.edu/pgc/download-results/ed/. Significant cell-types were identified using a Bonferroni p-value threshold of p<0.05.

### Cloning and DIG probe labeling

PCR with forward and reverse primers (see **Table 1**) was performed on mouse hypothalamic cDNA (isolated with miRNeasy Mini Kit, Qiagen, Hilden, Germany). PCR products were ligated into pGEMT.easy (Promega, Madison, WI, US) and sequenced with Sanger sequencing. PCR with SP6 and T7 primers was performed. cDNA probes were incubated for 2 hours at 37 degrees with Dig RNA labeling mix (11277073910, Roche, Basel, Switzerland) and SP6 or T7 RNA polymerase (RPOLSP6-RO and RPOLT7-RO, Roche).

**Table 1:**
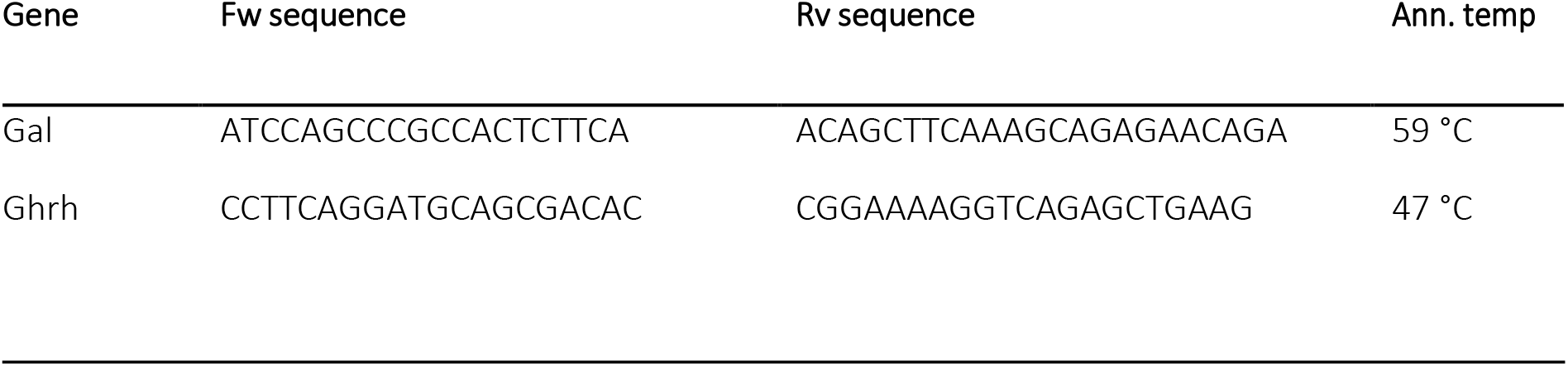
Primer sequences and annealing temperatures for PCR.

### Brain collection and sectioning for histology

LepR-Cre x L-tdTomato mice were perfused with PBS 1x followed by 4% paraformaldehyde (PFA) in PBS 1x. Brains were sliced coronally at 20μm thickness in a cryostat (Leica Biosystems, Wetzlar, Germany) with chamber temperature of 21 ± 3°C and object temperature of 19±3°C and sections were mounted directly on Superfrost glass (631-0108, VWR, Leuven, The Netherlands) in series of 10 per each brain and stored at −80°C.

### Fluorescent in situ hybridization and immunofluorescence for tdTomato

Slides were thawed at room temperature (RT) for 1 hour. Sections were incubated with 1,32% triethanolamine and 0,18% HCl for 10’ at RT and with hybridization mix (50% deionized formamide, 5x SSC buffer, 5x Denharts, 250 μg/ml tRNA baker’s yeast and 500 μg/ml Sonificated Salmon Sperm DNA) for 2 hours at RT. Sections were incubated with RNA probes (400 ng/mL hybridization mix) overnight at 68 degrees. Slides were transferred to 2x SSC at 68 degrees and then immediately to 0.2x SSC at 68 degrees for 2 hours. After that, sections were treated with 0,3% hydrogen peroxide in 1x TBS for 30’ at RT, blocked with TNB blocking buffer (from TSA Plus Cyanine 3 System, NEL744001KT, PerkinElmer, Waltham, MA, US) for 1 hr at RT and incubated with anti-DIG-POD (1:500, 11207733910, Roche) and rabbit anti-RFP (1:1000, 600-401-379, Rockland, Limerick, PA, US) in TNB blocking buffer overnight at 4 degrees. Sections were then incubated with Cyanine 3 Tyramide amplification reagent (1:50 in 1X amplification diluent) from TSA kit for 10-15’ at RT followed by incubation with goat anti-rabbit 568 (1:500, ab175471, Abcam, Cambridge, UK) in TNB blocking buffer and DAPI (1:1000 in 1x PBS). Between steps sections were washed 4×5’ with 1xTNT buffer (0,1M Tris-HCl, pH7.5, 0,15M NaCl, 0,05% Tween20). Sections were let to dry and covered with Fluorsave reagent (Calbiochem, San Diego, CA, US).

### Immunofluorescence

Slides were thawed at room temperature (RT) for 1 hour. Slides were incubated with blocking solution (5% normal goat serum (NGS), 0.5% Triton X-100 in 1x PBS) for 1 hour at room temperature (RT), followed by 2 hours incubation at RTwithRabbit anti-TH (Milipore, Ab152)1:500 diluted in carrier solution (5% NGS, 0,1% Triton X-100 in 1x PBS). Sections were then incubated with Alexa fluor 488 Goat a-rabbit (ab150077, Abcam) 1:1000 diluted in carrier solution for 2 hours at RT. Between all steps sections were washed 3 times for 5-10 minutes in PBS 1x. Sections were let to dry and covered with Fluorsave (Calbiochem, San Diego, CA, US).

### Imaging and Image analysis

10x magnification pictures were taken with an epi-fluorescent microscope (Zeiss Scope A1, ZEISS, Germany). Hypothalamic structures were defined based on the The Mouse Brain in Stereotaxic Coordinates, 3rd Edition^124^.

### Data and code sharing

The scRNAseq raw data is uploaded to GEO (#XXX) in the format of fastq files and a preprocessed dataset as a csv file. Data of Campbell et al was downloaded from GEO (GSE93374). All the code used for the analysis in the single cell data as well as the codes to reproduce the figures of the manuscript is deposited to Github (https://github.com/neuronur/scLepr).

---

**Figure S1.**
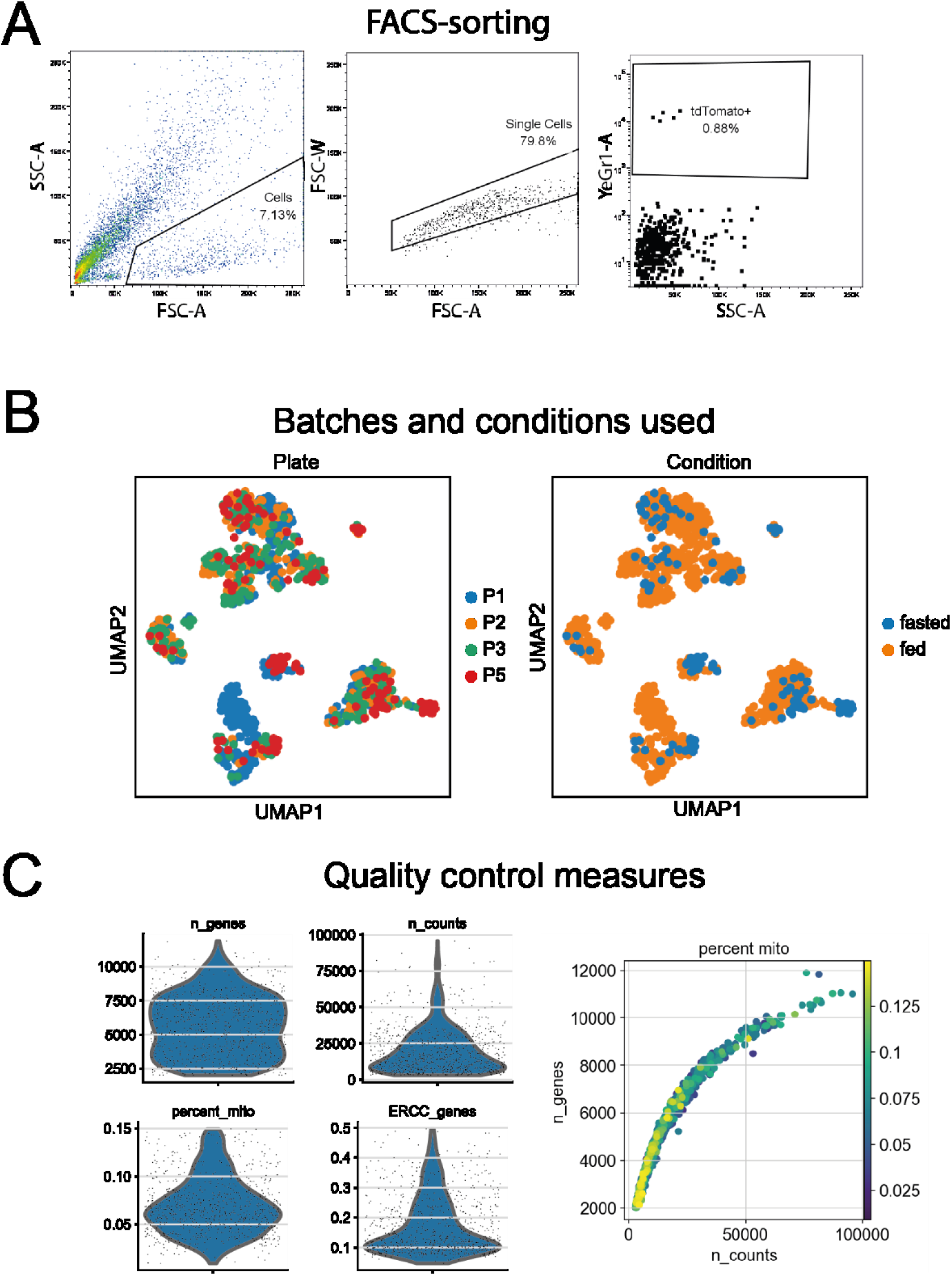
FACS-sorting, plate/batch and condition distribution and quality measures. **A** Plots representing the selection of cells to be sorted in 384-well plates, example from a single run. Left: 7,13% of the population was selected from the forward (FSC) and side scatter (SSC). Middle: 79,8% of cells were selected as singlets. Right: 0.88% of singlets were selected as tdTomato-positive based on fluorescence detected by the yellow laser. **B** UMAP plots showing distribution of plates (batches) and conditions **C** Plots representing quality measures for the number of genes, counts, percentage of mitochondrial genes and fraction of ERCC spike-in genes. The plot on the right show the correlation between the number of genes and counts as a measure of complexity. Cells are colored with the percentage of mitochondrial genes detected.

**Figure S2.**
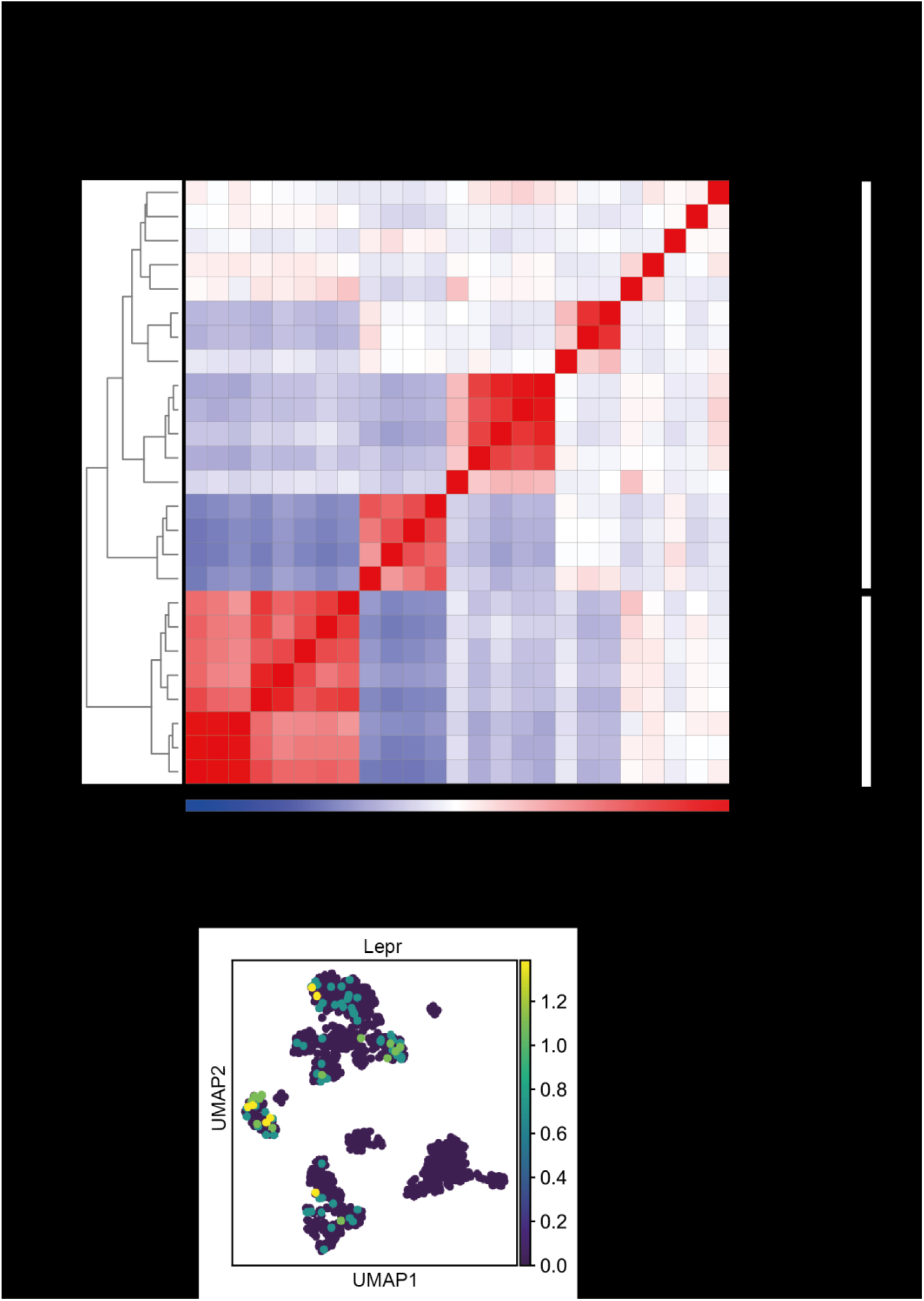
Hierarchical clustering of clusters and *Lepr* expression. **A** Heatmap showing Euclidean distances between clusters. The dendrogram depicts hierarchical clustering. **B** UMAP plot of Lepr gene expression. The expression level (normalized counts) is color-coded.

**Figure S3.**
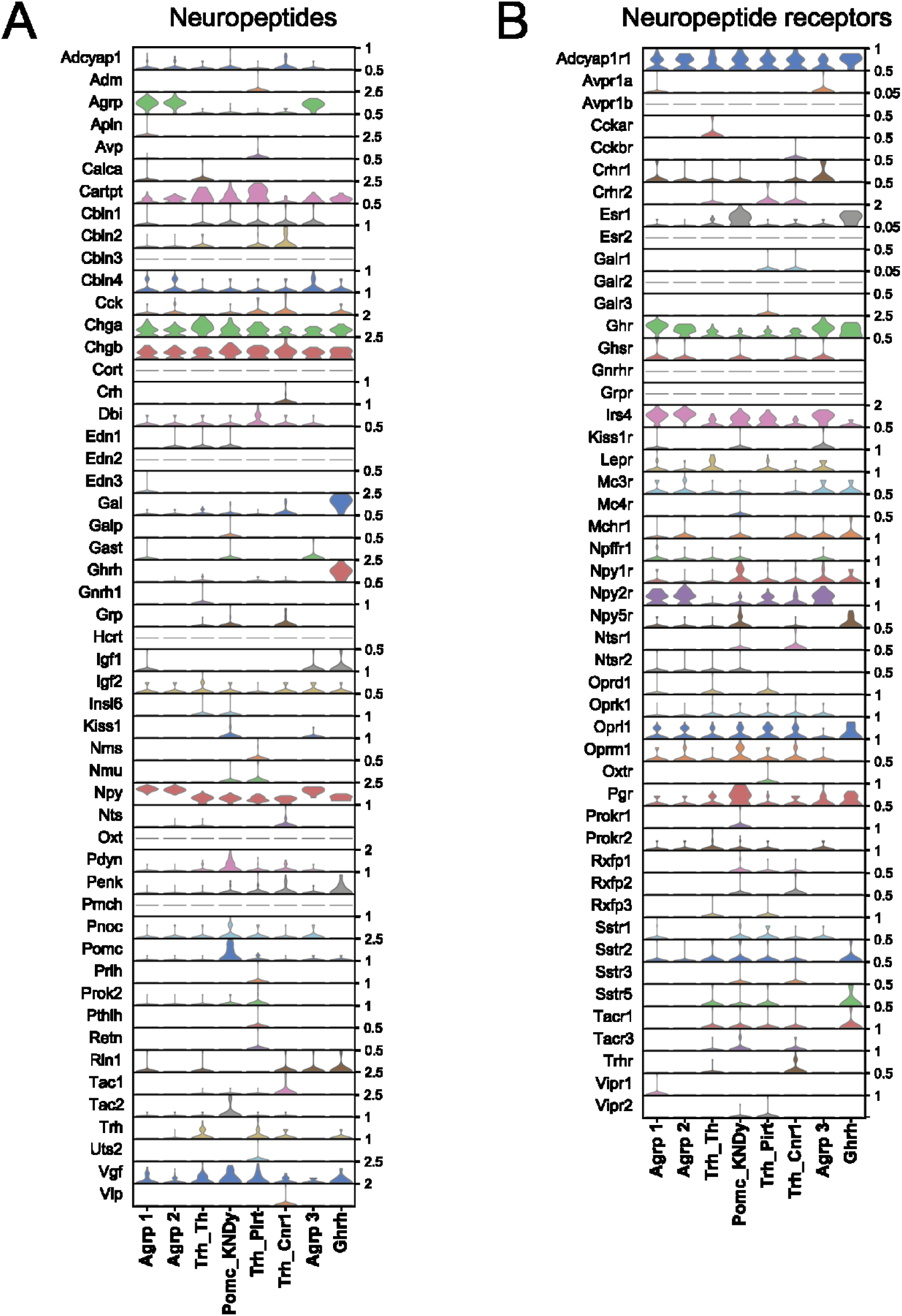
Neuropeptide and neuropeptide receptor expression in neuronal clusters of LepR neurons in the hypothalamus. **A-B** Violin plots of expression of neuropeptide (A) and neuropeptide receptor genes (B) in neuronal clusters. The maximum normalized count of each gene is presented on the right.

**Figure S4.**
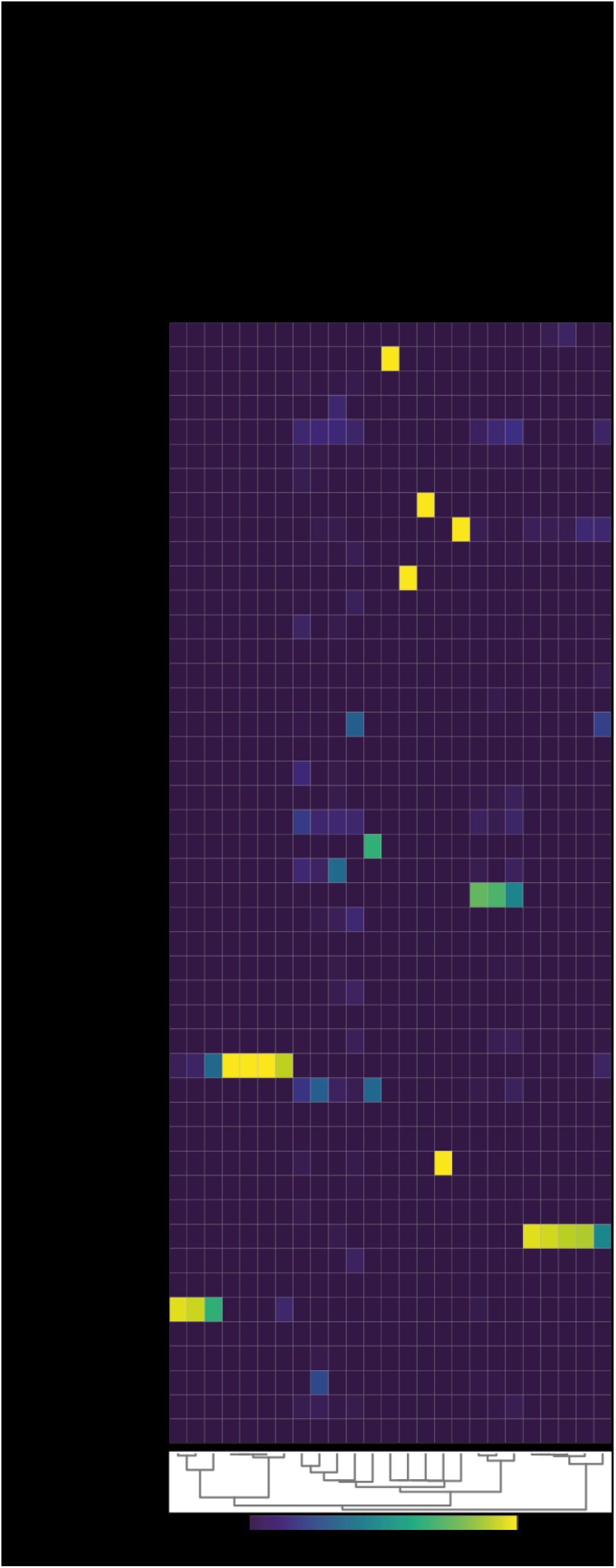
Comparison of Lepr cell types to the literature. Matrixplot showing the predicted probability of assignment of Lepr cell types identified in our study to published datasets using the SciBet pipeline. Chen et al identified cell types from the whole hypothalamus. Endo: Endothelial, Astro: Astrocytes, Tany: Tanycytes, VLMC: Vascular leptomeningeal cells, MM: Microglia and Macrophages, Oligo: Oligodendrocytes, OPCs: Oligodendrocyte progenitor cells.

**Figure S5** pySCENIC analysis for identification of GRNs

**A** tSNE plot representing distribution of conditions

**B** Dot-plot representing the 38 regulons identified to be active in the 25 clusters of LepR hypothalamic cells. Color-coding represents average regulon scores per cell and dot size the percentage of cells in which the regulon is active.

**Figure S6.**
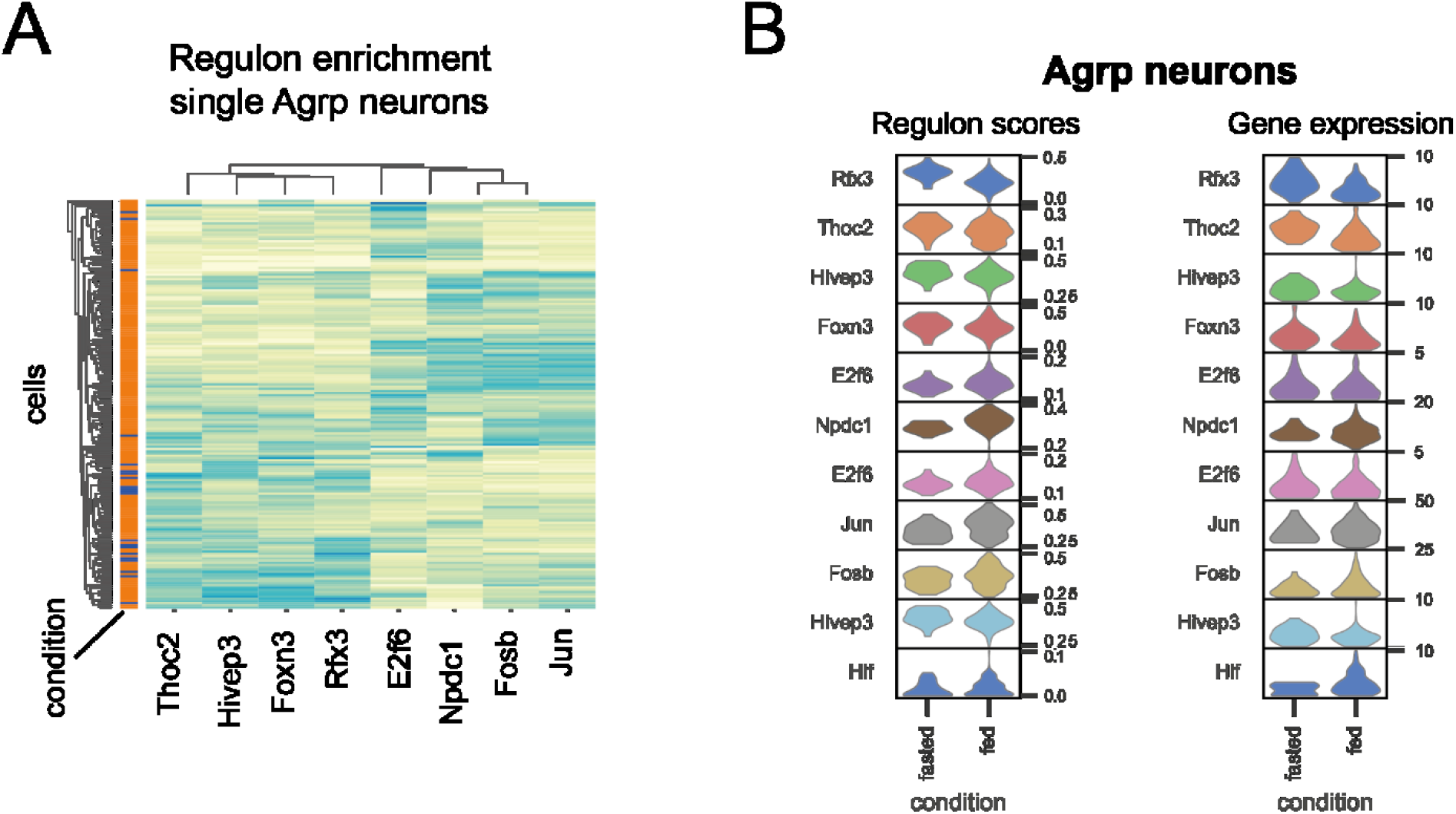
GRNs regulated by fasting in LepR Agrp neurons. **A** Heatmap representing regulon activity in individual LepR Agrp neurons from the fed and fasted condition. Color-coding represents regulon score. **B** Violin plots of regulon scores (numbers represent min and max) and gene expression levels (numbers represent max) in the fed and fasted condition

## Table legends

**Sup. Table 1** Differential expression analysis of genes in Agrp LepR cells of the fed and fasted condition. Mean: mean counts in fed/fasted, pct: fraction of cells expressing the gene, s: score, l: log2FoldChange, p: adjusted p value.

**Sup. Table 2** Target genes of each regulon identified to be differentially regulated upon fasting in Agrp LepR cells.

